# Stability of four carbapenem antibiotics in discs used for antimicrobial susceptibility testing

**DOI:** 10.1101/2024.06.17.599257

**Authors:** Selvi N. Shahab, Michiel L. Bexkens, Nikita Kempenaars, Amber Rijfkogel, Anis Karuniawati, Margreet C. Vos, Wil H.F. Goessens, Juliëtte A. Severin, the SAMPAN Consortium

**Affiliations:** Department of Medical Microbiology and Infectious Diseases, Erasmus MC University Medical Centre, Rotterdam, The Netherlands; Department of Clinical Microbiology, Faculty of Medicine Universitas Indonesia / Dr. Cipto Mangunkusumo General Hospital, Jakarta, Indonesia

**Keywords:** Carbapenems, drug stability, microbial sensitivity test

## Abstract

In low-to middle-income countries, microbiological laboratories often use disc diffusion for antimicrobial susceptibility testing (AST). Reliable AST of carbapenem antibiotics is crucial for treatment decisions and surveillance purposes. Transport and storage conditions of materials used for AST are critical and may be challenging in some settings, where temperature cannot always be controlled. This study aimed to test the stability of four carbapenems in discs for AST under unfavourable conditions, *i*.*e*., at room temperature and 35°C for up to 72 hours. Imipenem, meropenem, ertapenem, and doripenem discs from three brands, Oxoid, Becton Dickinson, and HiMedia, containing 10 μg of antibiotic were included. Discs were exposed to six unfavourable conditions and the recommended storage-condition as control. Subsequently, disc diffusion testing following the EUCAST guidelines was performed with four well-defined strains of *Escherichia coli* with different susceptibility profiles to carbapenems. The inhibition zone diameters were measured after 16-18 hours of incubation at 35±2°C. All experiments were executed in triplicate. In parallel, the carbapenems’ degradation was observed using a spectrophotometric method. Our study revealed that carbapenem discs were generally stable for AST although the concentration of most carbapenem antibiotics in discs decreased over time. Overall, imipenem (Oxoid and Becton Dickinson) discs were the most stable. Meropenem discs were less stable when exposed to 35°C than at room temperature. Concentrations of carbapenems in HiMedia discs were higher than those in Oxoid and Becton Dickinson. For carbapenem AST using disc diffusion in a rural area, we recommend using imipenem discs from Oxoid or Becton Dickinson.

**GRAPHICAL ABSTRACT:** 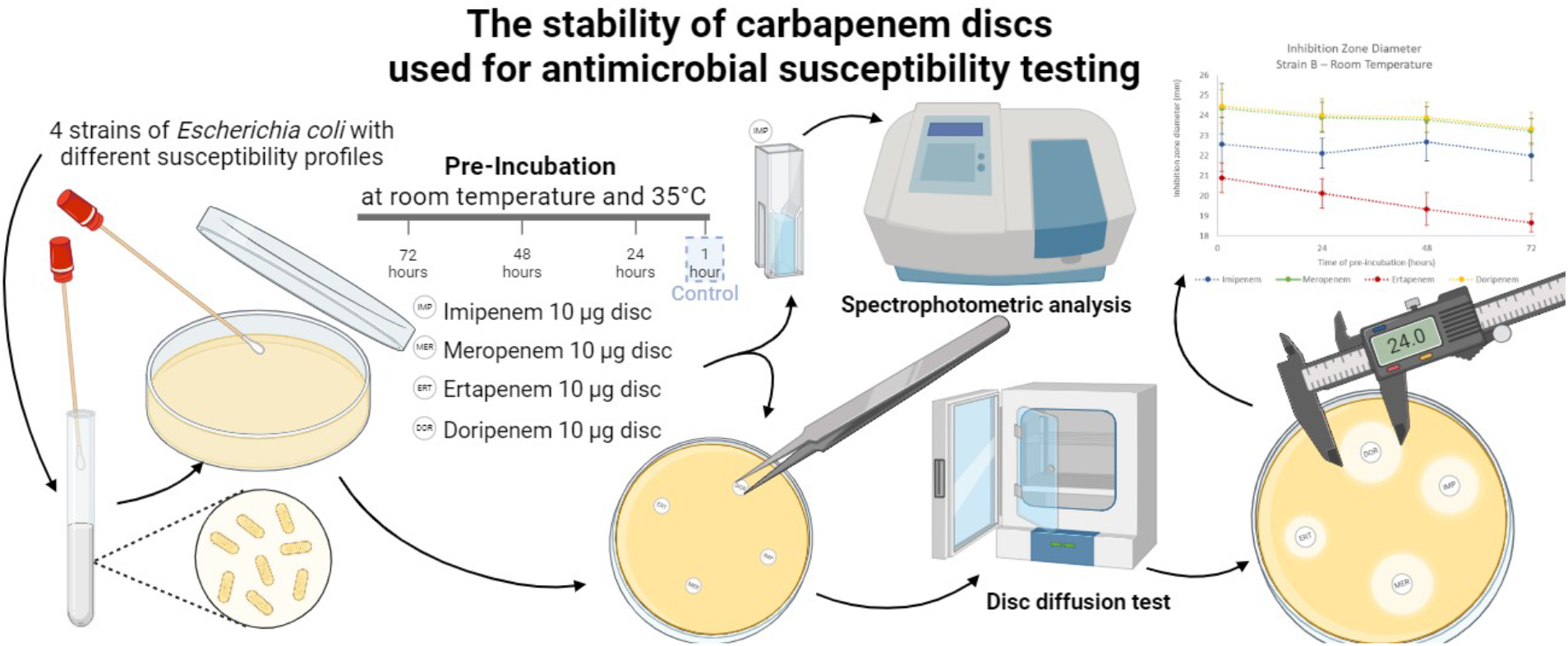

**HIGHLIGHTS:**

- In remote areas, transportation conditions of AST discs may be challenging
- Discs with four carbapenems from three brands were tested using six conditions
- Carbapenem discs were generally stable for AST
- In unfavourable conditions, concentrations in the discs degraded over time
- Imipenem (Oxoid and Becton Dickinson) discs were the most stable

## INTRODUCTION

Carbapenem antibiotics are beta-lactam antibiotics with a broad spectrum of activity that includes a wide range of multidrug-resistant (MDR) bacteria.[1] Moreover, carbapenems are considered to be the most reliable last-line treatment option for severe infections by MDR Gram-negative bacteria, especially extended-spectrum beta-lactamase-producing Enterobacterales.[2, 3] However, resistance to carbapenem antibiotics is on the rise worldwide and the World Health Organization (WHO) ranks carbapenem-resistant *Acinetobacter baumannii, Pseudomonas aeruginosa*, and Enterobacterales critical in the list of ‘priority pathogens’ for which action is needed.[4]

With the rise of carbapenem-resistant strains, it is of utmost importance that antimicrobial susceptibility testing (AST) for carbapenems is performed accurately in order to decide whether a carbapenem can be used as treatment for an infection and if additional infection prevention and control measures have to be taken.[5, 6] The disc diffusion test is one of the AST methods that is widely used around the globe, especially in areas with limited resources. This test is simple, cheap, and practical to perform. However, correct results will only be obtained when laboratories use materials of good quality, adhere strictly to the methodology and conduct rigorous quality control.[7] Any breach in this process may lead to incorrect results with major consequences such as failure of treatment.

One important aspect of the disc diffusion test is the antibiotic disc itself which can be purchased from several manufacturers. A previous study showed that the quality of discs from different manufacturers varied and only three of nine manufacturers maintained a high standard.[8] Manufacturers usually provide a guide for the storage and use of the discs and laboratories are responsible to adhere to these guidelines. However, laboratories are also dependent on reagents arriving in a good condition at their site. Maintaining the conditions during transport from the manufacturer to the laboratory may be a challenge, especially when the laboratory is remote, infrastructure is underdeveloped, or outdoor temperatures are high.[9] When antibiotic discs are exposed to unfavourable conditions, such as higher temperatures than recommended, antibiotics may degrade which will affect disc diffusion results, resulting in major errors, shifting the interpretation from susceptible to resistant. To the best of our knowledge, this has not been investigated in detail before.

This study aimed to investigate the stability of carbapenem discs under unfavourable conditions that may occur during transportation. The carbapenem discs were pre-incubated mimicking two unfavourable shipping temperatures, each for three durations. The discs were then used for performing the disc diffusion test of four different strains of *Escherichia coli* and the concentrations of carbapenem in the discs were measured using spectrophotometric testing. The findings of these experiments reveal the effects of unideal conditions on the carbapenem discs.

## MATERIALS AND METHODS

### Discs and Pre-incubation Conditions

Carbapenem discs containing 10 µg of the active ingredient (either imipenem, meropenem, ertapenem, or doripenem) were used from three manufacturers: Oxoid (Thermo Fisher Scientific; CT0455B, CT0774B, CT17161B, and CT1880B), Becton Dickinson (BBL^TM^; 231645, 231703, 232174, and 232219), and HiMedia (SD073, SD727, SD283, and SD280). These three brands were chosen because they are used in a global scale. Each disc had a diameter of 6 mm and was kept at 4°C temperature in dark conditions before the experiment as per the manufacturer’s instruction. Nearly before the experiment, the discs were exposed to six different unfavourable conditions and one hour at room temperature as a control. The durations for unfavourable pre-incubation conditions were 24, 48, and 72 hours at both room temperature and 35°C. Those conditions were considered to mimic the possible shipping period and temperature.

### Bacterial Strains

Four well-defined strains of *E. coli* with different susceptibility profiles to carbapenems were used. One strain (strain A) was the quality control (QC) strain ATCC 25922 and the others were taken from the Erasmus MC collection (Table 1). All strains were identified using matrix-assisted laser desorption/ionization time-of-flight mass spectrometry (MALDI-TOF MS; Bruker Daltonics GmbH, Bremen, Germany). Initial antimicrobial susceptibility testing was performed using VITEK2® (bioMérieux, Inc., Marcy-l’Etoile, France). MICs were further determined with broth microdilution (Trek Diagnostic Systems, Thermo Fisher Scientific, Franklin, USA). MICs were interpreted according to EUCAST Breakpoint Tables v. 10.0, 2020 for imipenem, meropenem, and ertapenem and v. 8.1, 2018 for doripenem.[10, 11] Carbapenem resistance genes were determined using PCR, when applicable.[12]

**Table 1.**
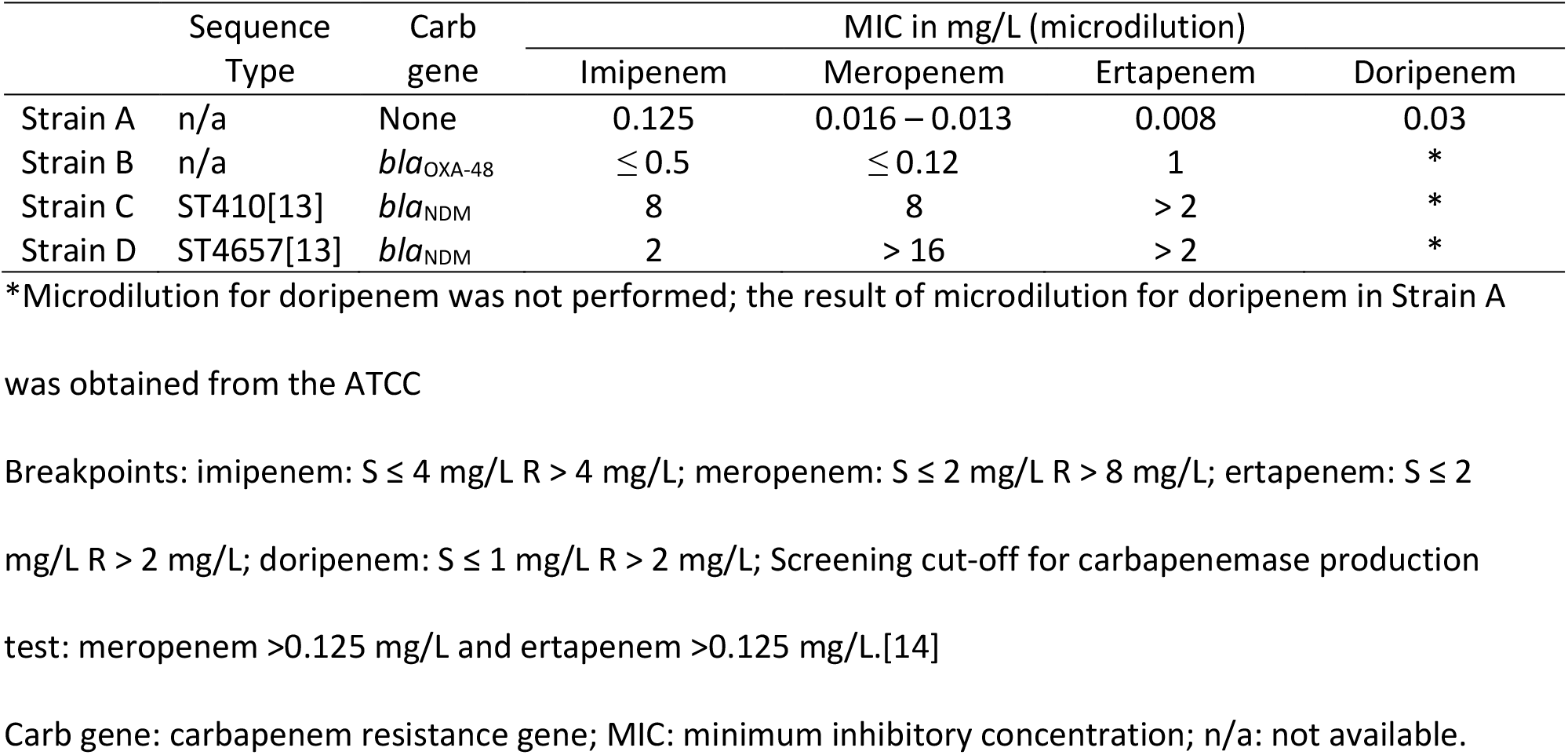
Strains of Escherichia coli used in this research

### Disc Diffusion Test

A standard Kirby-Bauer disc diffusion test was used following the EUCAST guidelines using the pre-incubated carbapenem discs to test the susceptibility patterns of four different strains of *E. coli* following previously described method.[15] Before the procedure, all strains were freshly cultured on blood agar plates (Becton Dickinson) for 18-24 hours. Each strain was inoculated on twenty-one separate different Mueller-Hinton II agar plates (Becton Dickinson). To check for purity, the suspensions of the strains used for the test were cultured on blood agar plate.

On the wet agar plates, the pre-incubated discs were applied using sterile forceps. Gentle pressure was made to the disc to make sure complete contact of the disc. On each Mueller-Hinton II agar plate, four different carbapenem discs, each from its unfavourable pre-incubation conditions, were placed with more than 24 mm distance. The discs were never relocated after they contacted the agar plate surface. Within 15 minutes of the discs’ application, the plates were incubated at 35°C.

After 16-18 hours of incubation, the inhibition zone diameter was measured using a black non-reflecting surface as the background. The measurement was performed using a standardized sliding calliper to the nearest millimetre, including the disc diameter. Zone inhibition diameters in strain A were compared to the EUCAST QC for disc diffusion version 8.0.[16] All the inhibition zone diameters were recorded and analysed using Microsoft Excel and International Business Machines (IBM) Statistical Package for Social Sciences (SPSS) version 26. Each experiment was executed three times.

### Spectrophotometric Analysis

In addition to disc diffusion testing, spectrophotometric analysis was used to observe the degradation of the carbapenem antibiotic in the discs. The degradation of imipenem, meropenem, ertapenem, and doripenem in the discs from the above mentioned three manufacturers was measured. The degradation was observed as the concentration of carbapenem after separate discs were exposed to the six conditions described previously and one control condition. The concentration of the carbapenem inside the discs after 1-hour in room temperature was recorded as the control (100%). After the pre-incubation period, each disc was placed in 150 µL of sterile aquadest, of which 10 µL was placed in the 96-well microtiter plate and measured at 300 nm. A standard curve for each antibiotic was made using the antibiotic powder (imipenem monohydrate ≥98%, I0160, Sigma-Aldrich; meropenem trihydrate ≥98%, M2574, Sigma-Aldrich; ertapenem sodium ≥90%, SML1238, Sigma-Aldrich; doripenem monohydrate ≥98%, 32138, Supelco) which was used to measure the amount of carbapenem released from the disc. The experiment was performed in triplicate, the mean of the concentrations from all three experiments was recorded as the result. The results were analysed and illustrated using Microsoft Excel.

## RESULTS

### Disc Diffusion Test

Inhibition zone diameters per condition, per strain, and per manufacturer are shown in Figures 1-3. In strain A, diameter zones of imipenem, meropenem, and ertapenem from Oxoid and Becton Dickinson after all pre-incubation conditions were in the QC range and ±2 mm from the QC target. While doripenem from Oxoid created inhibition zone diameters with minor deviation (±1 mm from the QC target), doripenem from Becton Dickinson created >2 mm smaller inhibition zone diameter than QC target but still within the QC range. Without unfavorable pre-incubation conditions, imipenem and doripenem discs from HiMedia created larger zones than the QC target (imipenem 29 mm and doripenem 31 mm) and even larger than the QC range (imipenem 26-32 mm and doripenem 27-35 mm). Meropenem and ertapenem discs from HiMedia also created larger zones than the QC target (meropenem 31-32 mm and ertapenem 32-33 mm) but still in the QC range (meropenem 28-35 mm and ertapenem 29-36 mm). The longer pre-incubation period, the smaller inhibition zone created by imipenem discs from HiMedia in strain A (from 34.78±0.79 mm to 27.11±1.44 mm at room temperature and to 26.22±1.23 mm at 35°C).

**Figure 1.**
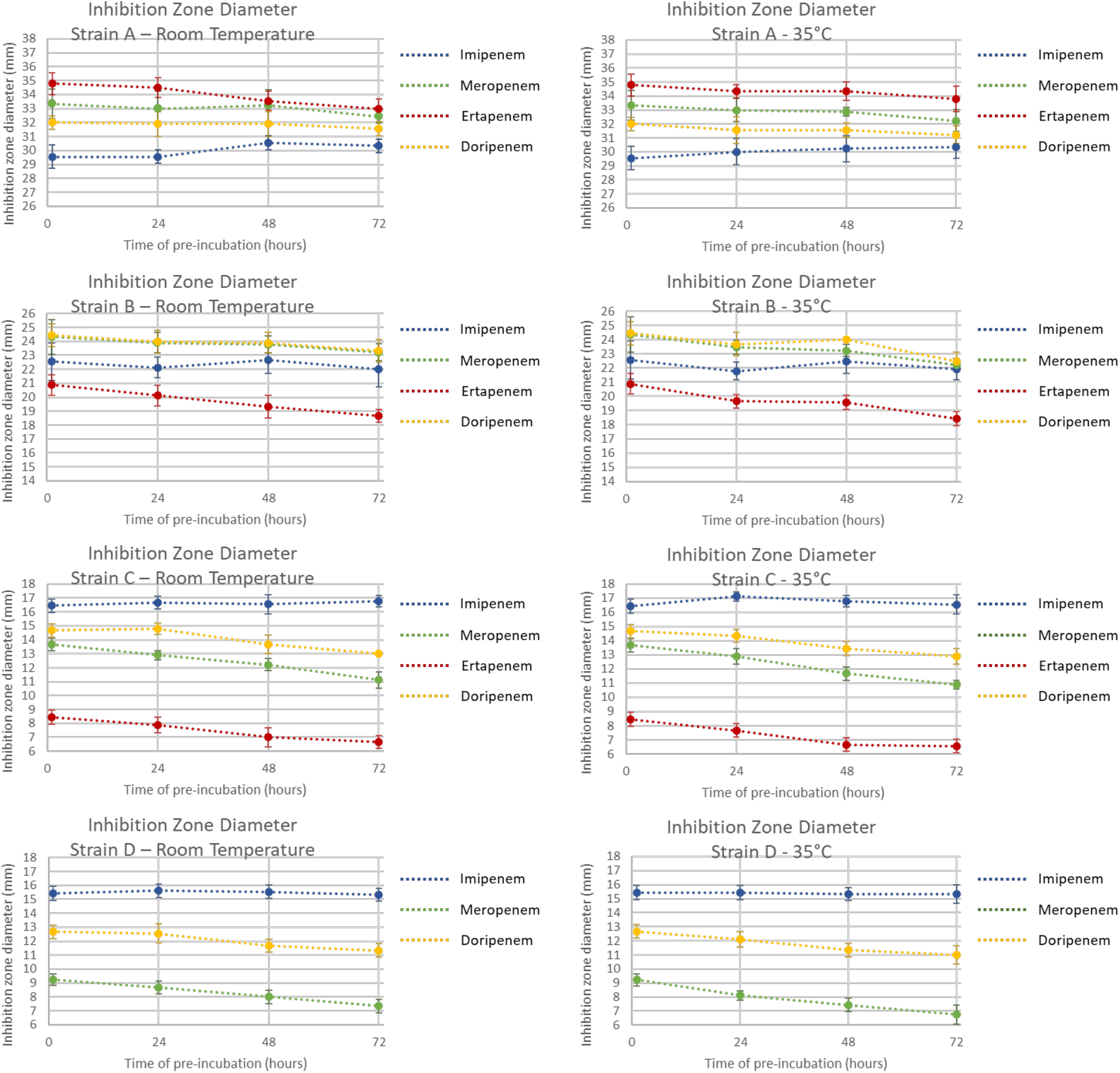
Comparison of different carbapenem discs from Oxoid in creating inhibition zone diameters by using disc diffusion test after pre-incubation at room temperature and 35°C. No inhibition zone observed for ertapenem disc in strain D.

**Figure 2.**
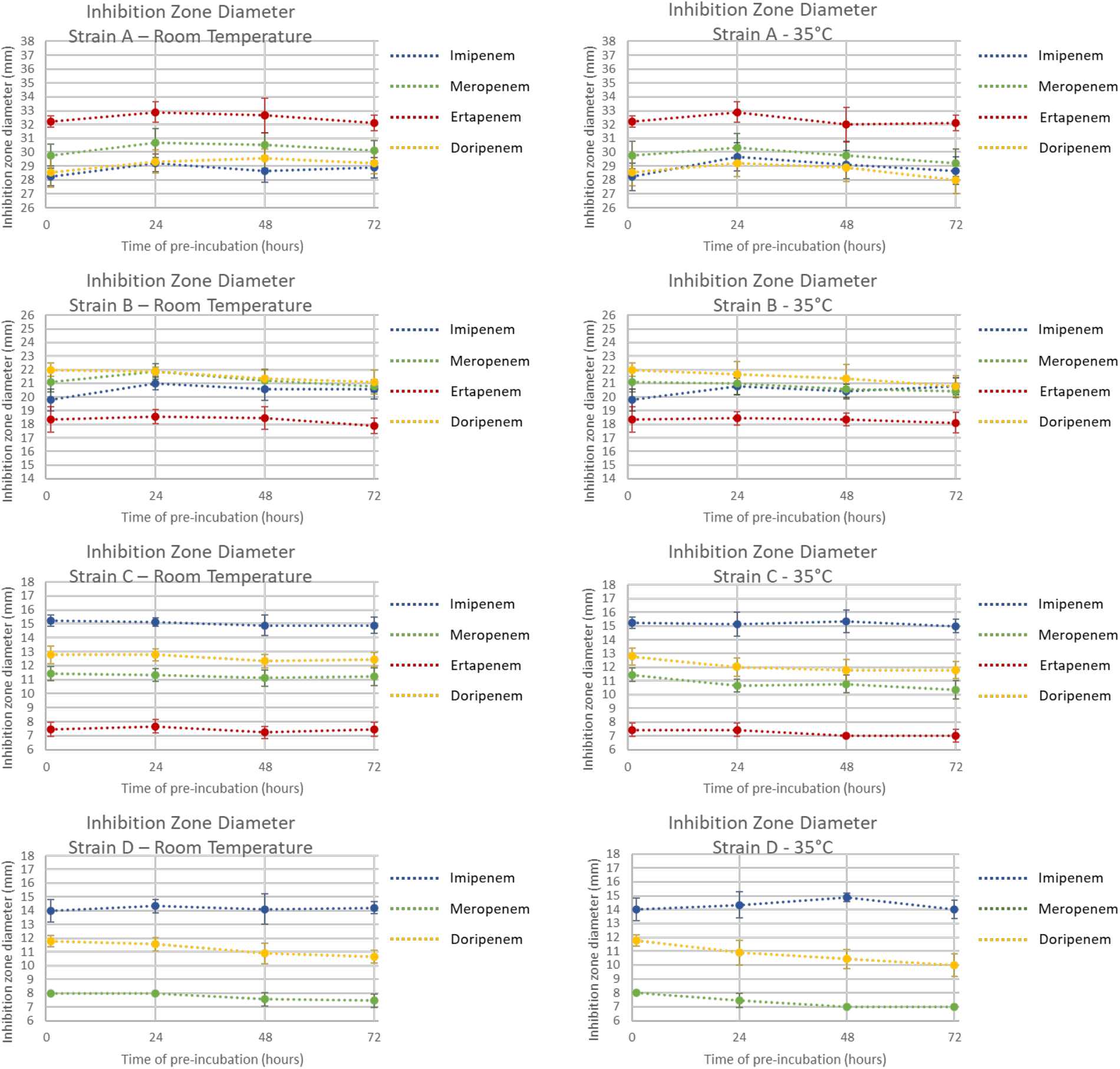
Comparison of different carbapenem discs from Becton Dickinson in creating inhibition zone diameters by using disc diffusion test after pre-incubation at room temperature and 35°C. No inhibition zone observed for ertapenem disc in strain D.

Among Oxoid discs, imipenem produced the most consistent diameters in all strains with <1 mm drops of zone diameter after the longest pre-incubation period. Meropenem, ertapenem, and doripenem discs consistently produced smaller inhibition zone diameters after the longer pre-incubation periods when used for strain B-D (Figure 1) with ertapenem having the largest fall in inhibition zone in strain B (from 20.89±0.74 mm to 18.67±0.47 mm at room temperature and to 18.44±0.50 mm at 35°C; all interpreted as resistant while the MIC data showed susceptibility) and meropenem having the largest fall in inhibition zone in strain C and D (strain C: from 13.67±0.47 mm to 11.11±0.57 mm at room temperature and to 10.89±0.31 mm at 35°C; strain D: from 9.22±0.42 mm to 7.33±0.47 mm at room temperature and to 6.75±0.66 mm at 35°C). Even with the smaller inhibition zone, the category interpretations did not change and still corresponded to the interpretation based on the MIC of the strains.

All carbapenem discs from Becton Dickinson created relatively stable zone diameters in all strains, showing inhibition zone diameters differences of ≤ 1 mm, except for meropenem in strain C at 35°C pre-incubation (from 11.44±0.50 mm to 10.33±0.67 mm) and doripenem in strain B at 35°C pre-incubation (from 22.00±0.47 mm to 20.78±0.79 mm) and strain D (from 11.78±0.42 to 10.67±0.47 mm at room temperature and to 10.00±0.82 at 35°C). The changes in inhibition zone diameters did not change the category interpretations. All inhibition zones formed for strain B using ertapenem discs, in the unfavourable conditions and the control condition, were in the resistance range, while the MIC as tested with microdilution was in the susceptible category. Across all strains, the imipenem disc provided the most stable inhibition zones.

Imipenem discs from HiMedia showed decreased zone diameters in all strains (Figure 3). The decreased zone diameters did not change the category interpretation in strain A (susceptible), but it changed in the other strains (strain B: from susceptible to intermediate while the MIC of the strain was in the susceptible range; strain C and D: from intermediate to resistant while the MIC of the strain was in the resistance and susceptible range, respectively). Although doripenem discs created the most consistent diameters, they always formed larger diameters compared to the other brands in all strains. A similar finding was observed for ertapenem discs. For instance, in strain D, while no inhibition zone was formed for Oxoid and Becton Dickinson discs, HiMedia ertapenem discs still formed inhibition zones despite the fact of being smaller after longer pre-incubation period (Figures 1-3).

**Figure 3.**
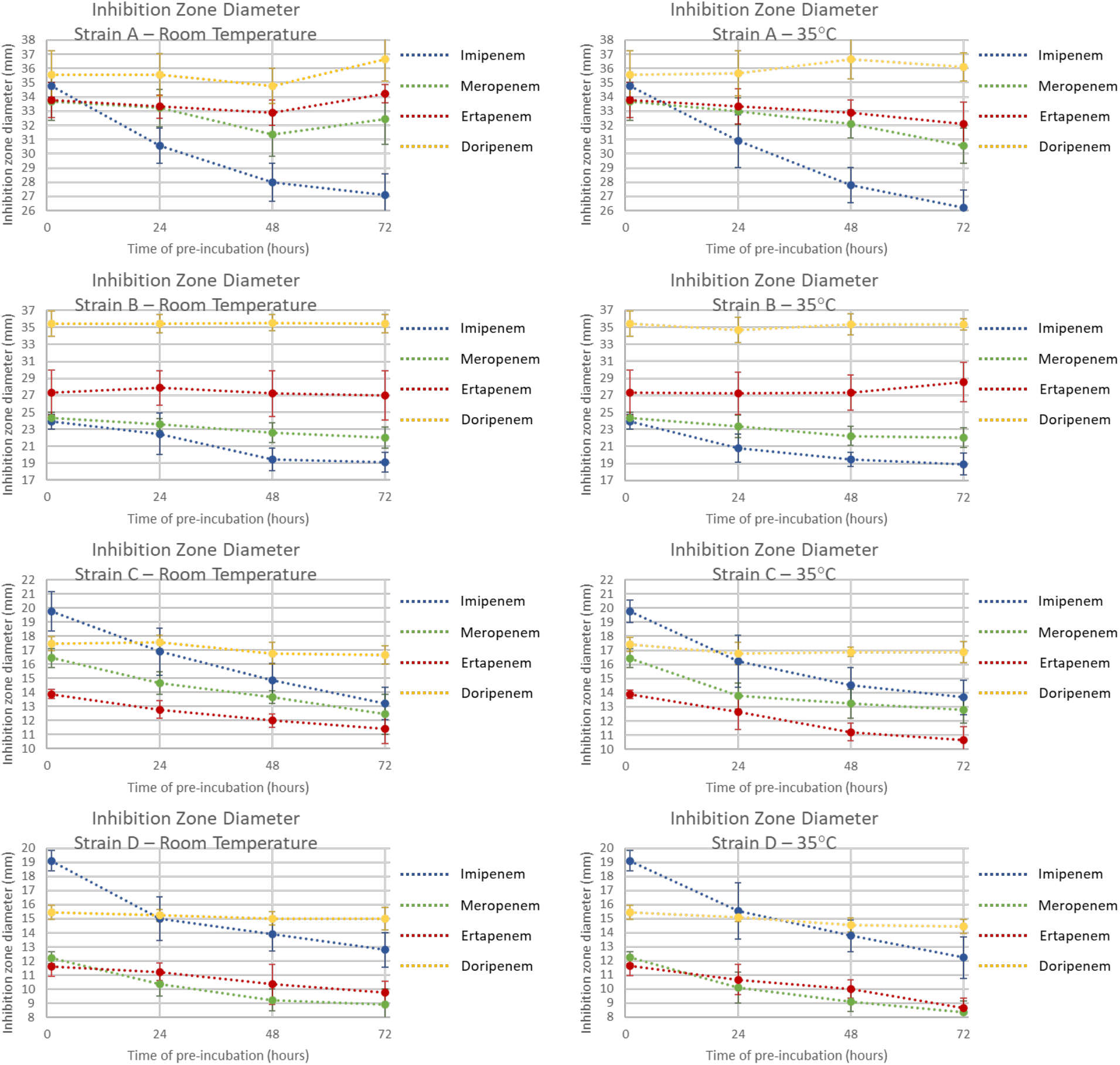
Comparison of different carbapenem discs from HiMedia in creating inhibition zone diameters by using disc diffusion test after pre-incubation at room temperature and 35°C.

### Spectrophotometric Analysis

Table 2 shows the control concentration used as baseline to measure the percentage of the concentration of the carbapenem. Concentrations of carbapenems in HiMedia discs were higher than those in Oxoid and Becton Dickinson. In general, most carbapenem antibiotics in the discs degraded over time based on the spectrophotometric analysis results (Figure 4). The degradation was more obvious when the discs were pre-incubated at 35°C. Imipenem discs from Oxoid had stable concentrations. While imipenem was only degraded for <20% in Becton Dickinson discs, the concentration of imipenem in HiMedia discs was degraded for >40% after 72 hours. The spectrophotometric analysis also revealed that meropenem from all brands degraded more at 35°C than at room temperature. When exposed to 35°C for 72 hours, the meropenem concentration was degraded for 69%, 27% and 52%, for Oxoid, Becton Dickinson, and HiMedia discs, respectively. While showing <10% degradation in Becton Dickinson and HiMedia discs, doripenem in Oxoid disc showed 26% and 73% degradation when exposed to room temperature and 35°C for 72 hours. Ertapenem concentration in Oxoid discs was also decreased by 14% and 62% when exposed to room temperature and 35°C for 72 hours while it remained stable in Becton Dickinson and HiMedia discs.

**Figure 4.**
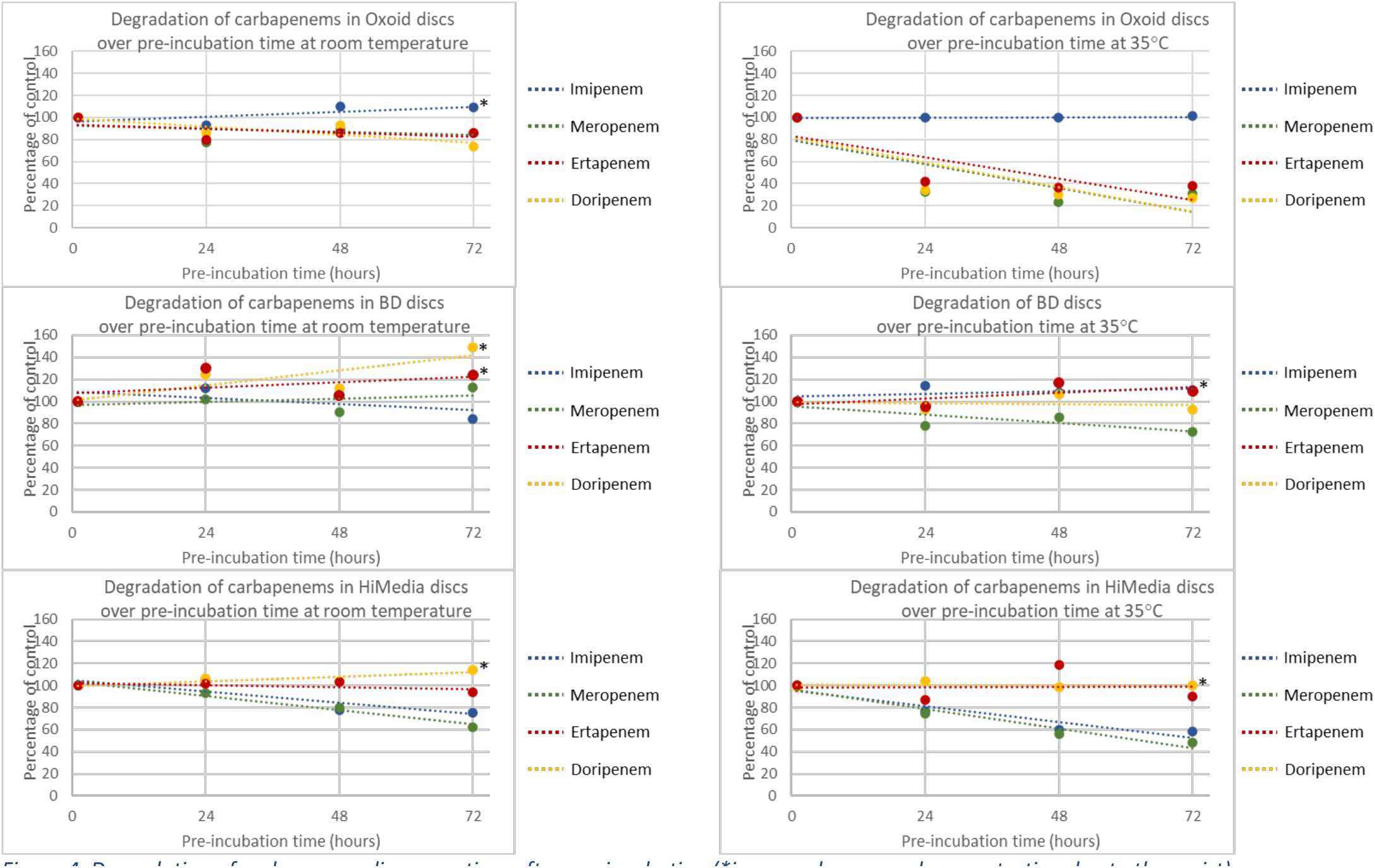
Degradation of carbapenem discs over time after pre-incubation (^*^increased measured concentration due to the moist)

## DISCUSSION

Our study revealed that carbapenem discs were generally stable for AST even after unfavourable pre-incubation conditions except for imipenem from HiMedia where the decreased inhibition zone diameters caused changes in category interpretation. This demonstrates that the use of disc diffusion test as AST for carbapenem antibiotic is a reliable method, even when it is performed in a remote area with a longer period of shipping. From this study, imipenem (Oxoid and Becton Dickinson), and doripenem (HiMedia) were the most stable carbapenem antibiotics in disc for disc diffusion test. However, doripenem from HiMedia formed larger diameters than the QC target and range. This finding was similar with a previous study comparing antimicrobial discs from nine manufacturers where inhibition zone diameters from HiMedia discs were outside the QC target.[8] The spectrophotometric analysis also confirmed the stability of imipenem concentration in Oxoid and Becton Dickinson discs. Moreover, the spectrophotometric analysis showed that concentrations of carbapenems in HiMedia discs were higher than those in Oxoid and Becton Dickinson. This explains why the inhibition zone diameters created by HiMedia discs in strain A were larger than QC target. Inconsistent results were also shown for strain B, a *bla*OXA-48-positive *E. coli*, for ertapenem (Oxoid and Becton Dickinson), where zone diameters were in the resistance range, while the MIC (1 mg/L) was in the susceptible category. However, this MIC reflects an indication to screen an isolate for carbapenemase production.[14]

Imipenem is an older carbapenem that is relatively more unstable than other carbapenems inside the body due to renal dehydropeptidase-I (DHP-I). For that, imipenem is used in combination with cilastatin, a DHP-I inhibitor.[17] In this study, where the activity of carbapenems was tested *in vitro*, with no DHP-I, imipenem (Oxoid and Becton Dickinson) showed good stability in all strains. Imipenem was mentioned as the indicator for detection of KPC carbapenemase-producing Enterobacterales in clinical practice.[18]

Meropenem is often used as the screening for carbapenem-resistant Enterobacterales and for detection of carbapenemase-producing Enterobacterales.[8] Nonetheless, our study showed that meropenem was less stable at a higher temperature and that meropenem was even the least stable carbapenem at 35°C based on the disc diffusion test (Oxoid and HiMedia) and the spectrophotometric analysis (Oxoid, Becton Dickinson and HiMedia). A previous study showed that meropenem as a drug in bone cement was relatively stable even after heat exposure.[19] The possible reason is that the study only exposed the drug for a maximum of 120 minutes. Our study corresponds to another previous study that showed that meropenem in normal saline solution was unstable at 37°C and that there was improved stability when the temperature was kept at below 25°C.[20] There is, however, some difficulty in controlling the temperature to below 25°C in tropical countries and even more in the shipping process unless more expensive measures are taken.[9]

Ertapenem and doripenem are relatively new carbapenems with promising outcomes of therapy.[21-23] Nevertheless, the size of inhibition zone formed by both ertapenem and doripenem discs had some differences when compared from one brand to the other. The novelty of the antibiotics might cause the standardization still needs to be improved. In addition, ertapenem was the least stable carbapenem at room temperature, in line with the previous study where ertapenem as drug was degraded to a lower concentration (below 90%) already after an hour in room temperature.[24] Another study suggested that ertapenem was only stable for about 5.5 hours in room temperature.[25] On the other hand, this study confirmed a previous study where doripenem mentioned to be more stable than meropenem.[26, 27] Some studies suggested doripenem as a drug was only stable for 12-16 hours at room temperature.[28, 29] Different from that data, this study showed doripenem disc was relatively stable even after more than 16 hours, especially in the experiments using the QC strain. However, when being used for the resistant strains, the stability was less than imipenem (Oxoid and Becton Dickinson) although doripenem from HiMedia showed otherwise. In that case, the high concentration of doripenem inside HiMedia disc should be considered.

To the best of our knowledge, no study has been published with experiments on the topic of the stability of carbapenem discs for AST. Changes of zone diameters formed for different carbapenem discs after different pre-incubation conditions were revealed. Our study has some limitations. First, our study was limited to carbapenem discs from three of nine manufacturers providing discs for antimicrobial susceptibility test. The aim was to compare every carbapenem to the others within each brand. Also, we choose three brands that are important on a global scale. Second, we only included discs with four carbapenem antibiotics. Other carbapenems, such as panipenem and biapenem, are on the market, and AST with disc diffusion could be applied for some of these.[30, 31] Third, although we observed that imipenem is the most stable carbapenem antibiotic, it should be noted that we did not look at the best indicator for screening for a carbapenemase. When using imipenem or meropenem, it is known that Enterobacterales with *bla*OXA-48 may be missed. Additionally, for some Enterobacterales (*Morganella morganii, Providencia* spp, *Proteus* spp, and *Serratia* spp), imipenem is also not recommended as indicator, due to the lower intrinsically activity towards these species.

## CONCLUSIONS

In conclusion, the exposure of unfavourable temperatures during the shipping for up to 72 hours will only slightly change the inhibition zone diameter, although differences between brands were observed. For carbapenem resistance screening using the disc diffusion test in a rural area with high risk on long periods of high temperature (above 4°C recommended temperature from the manufacturers), we recommend using imipenem as the carbapenem of choice, preferably from brand Oxoid or Becton Dickinson, as it was the most stable one.

## ACKNOWLEDGEMENTS

We are grateful to all members of the SAMPAN Consortium for their input: Anniek de Jong (Deltares, Delft, the Netherlands), Heike Schmitt (National Institute of Public Health and the Environment, Bilthoven, the Netherlands), Maurizio Sanguinetti (Fondazione Policlinico Universitario “A. Gemelli”, Rome, Italy), Nicola Petrosilli (National Institute for Infectious Diseases “L. Spallanzani”, Rome, Italy), Roger C. Lévesque (U. Laval Integrative Systems Biology Institute, Québec, Canada).

## CONTRIBUTIONS

J.A.S., S.N.S. and M.L.B. contributed to the study concept and design; S.N.S., N.K. and A.R. performed disc diffusion and spectrophotometric test. S.N.S., M.L.B. and J.A.S. analysed and interpreted the data.

S.N.S. wrote the initial draft of the manuscript. J.A.S., M.L.B., N.K., A.M., A.K., M.C.V. and W.H.F.G gave valuable input to the manuscript. All authors read, reviewed, and approved the final article.

## REFERENCES

[1] Nicolau DP. Carbapenems: a potent class of antibiotics. Expert Opin Pharmacother. 2008;9:23–37.

[2] Bouza E. The role of new carbapenem combinations in the treatment of multidrug-resistant Gram-negative infections. Journal of Antimicrobial Chemotherapy. 2021;76:iv38–iv45.

[3] Baughman RP. The use of carbapenems in the treatment of serious infections. J Intensive Care Med. 2009;24:230–41.

[4] World Health Organization. WHO publishes list of bacteria for which new antibiotics are urgently needed. 2017.

[5] Lee M, Chung HS. Different antimicrobial susceptibility testing methods to detect ertapenem resistance in Enterobacteriaceae: VITEK2, MicroScan, Etest, disk diffusion, and broth microdilution. J Microbiol Methods. 2015;112:87–91.

[6] van Dijk MD, Voor In ‘t Holt AF, Alp E, Hell M, Petrosillo N, Presterl E, et al. Infection prevention and control policies in hospitals and prevalence of highly resistant microorganisms: an international comparative study. Antimicrob Resist Infect Control. 2022;11:152.

[7] Åhman J, Matuschek E, Kahlmeter G. EUCAST evaluation of 21 brands of Mueller-Hinton dehydrated media for disc diffusion testing. Clin Microbiol Infect. 2020;26:1412.e1-.e5.

[8] Åhman J, Matuschek E, Kahlmeter G. The quality of antimicrobial discs from nine manufacturers-EUCAST evaluations in 2014 and 2017. Clin Microbiol Infect. 2019;25:346–52.

[9] Lowe DE, Pellegrini G, LeMasters E, Carter AJ, Weiner ZP. Analysis and modeling of coolants and coolers for specimen transportation. PLoS One. 2020;15:e0231093.

[10] EUCAST. Breakpoint tables for interpretation of MICs and zone diameters, version 10.0, 2020.

[11] EUCAST. Breakpoint tables for interpretation of MICs and zone diameters, version 8.1, 2018.

[12] Saharman YR, Karuniawati A, Sedono R, Aditianingsih D, Goessens WHF, Klaassen CHW, et al. Clinical impact of endemic NDM-producing Klebsiella pneumoniae in intensive care units of the national referral hospital in Jakarta, Indonesia. Antimicrob Resist Infect Control. 2020;9:61.

[13] van der Zwaluw K, Witteveen S, Wielders L, van Santen M, Landman F, de Haan A, et al. Molecular characteristics of carbapenemase-producing Enterobacterales in the Netherlands; results of the 2014-2018 national laboratory surveillance. Clin Microbiol Infect. 2020;26:1412.e7-.e12.

[14] EUCAST. EUCAST guidelines for detection of resistance mechanisms and specific resistances of clinical and/or epidemiological importance, version 2, 2017.

[15] Matuschek E, Brown DF, Kahlmeter G. Development of the EUCAST disk diffusion antimicrobial susceptibility testing method and its implementation in routine microbiology laboratories. Clin Microbiol Infect. 2014;20:O255–66.

[16] EUCAST. Routine and extended internal quality control for MIC determination and disk diffusion as recommended by EUCAST version 8.0. 2018.

[17] Zhanel GG, Wiebe R, Dilay L, Thomson K, Rubinstein E, Hoban DJ, et al. Comparative review of the carbapenems. Drugs. 2007;67:1027–52.

[18] Benenson S, Temper V, Cohen MJ, Schwartz C, Hidalgo-Grass C, Block C. Imipenem disc for detection of KPC carbapenemase-producing Enterobacteriaceae in clinical practice. J Clin Microbiol. 2011;49:1617–20.

[19] Schmid M, Steiner O, Fasshold L, Goessler W, Holl AM, Kühn KD. The stability of carbapenems before and after admixture to PMMA-cement used for replacement surgery caused by Gram-negative bacteria. Eur J Med Res. 2020;25:34.

[20] Jaruratanasirikul S, Sriwiriyajan S. Stability of meropenem in normal saline solution after storage at room temperature. Southeast Asian J Trop Med Public Health. 2003;34:627–9.

[21] Jones RN, Huynh HK, Biedenbach DJ, Fritsche TR, Sader HS. Doripenem (S-4661), a novel carbapenem: comparative activity against contemporary pathogens including bactericidal action and preliminary in vitro methods evaluations. J Antimicrob Chemother. 2004;54:144–54.

[22] Dedhia HV, McKnight R. Doripenem: position in clinical practice. Expert Rev Anti Infect Ther. 2009;7:507–14.

[23] Wexler HM. In vitro activity of ertapenem: review of recent studies. J Antimicrob Chemother. 2004;53 Suppl 2:ii11–21.

[24] Kuti JL, Nicolau DP. Stability of Ertapenem 100 mg/mL at Room Temperature. Can J Hosp Pharm. 2016;69:256–9.

[25] Walker SE, Law S, Perks W, Iazzetta J. Stability of Ertapenem 100 mg/mL in Manufacturer’s Glass Vials or Syringes at 4°C and 23°C. Can J Hosp Pharm. 2015;68:121–6.

[26] Berthoin K, Le Duff CS, Marchand-Brynaert J, Carryn S, Tulkens PM. Stability of meropenem and doripenem solutions for administration by continuous infusion. J Antimicrob Chemother. 2010;65:1073–5.

[27] Jones RN, Sader HS, Fritsche TR. Comparative activity of doripenem and three other carbapenems tested against Gram-negative bacilli with various beta-lactamase resistance mechanisms. Diagn Microbiol Infect Dis. 2005;52:71–4.

[28] Crandon JL, Sutherland C, Nicolau DP. Stability of doripenem in polyvinyl chloride bags and elastomeric pumps. Am J Health Syst Pharm. 2010;67:1539–44.

[29] Psathas PA, Kuzmission A, Ikeda K, Yasuo S. Stability of doripenem in vitro in representative infusion solutions and infusion bags. Clin Ther. 2008;30:2075–87.

[30] Kimura K, Wachino J, Kurokawa H, Suzuki S, Yamane K, Shibata N, et al. Practical disk diffusion test for detecting group B streptococcus with reduced penicillin susceptibility. J Clin Microbiol. 2009;47:4154–7.

[31] Wang X, Zhang X, Zong Z, Yu R, Lv X, Xin J, et al. Biapenem versus meropenem in the treatment of bacterial infections: a multicenter, randomized, controlled clinical trial. Indian J Med Res. 2013;138:995–1002.

